# Identification of surface colors across environmental illuminations

**DOI:** 10.64898/2026.10.03.756408

**Authors:** Takuma Morimoto, Hannah E. Smithson

## Abstract

When objects are embedded in a complex lighting environment, the illumination they receive depends on their geometric relationship to the environment. We measured performance in an odd-one-out color constancy task, with objects rendered in complex lighting environments. Observers identified the odd object among four objects, with two rendered under each of two lighting environments, sampled from a wide range of object orientations and camera positions. Constancy was high. The presence of a surrounding context increased accuracy. Gloss provided no measurable benefit. We also tested a condition in which the global color impression provided misleading information about the lighting environment, but this impaired performance only slightly. These findings suggest that human color constancy remains robust even under complex lighting environments, contrary to predictions of many simple models of ‘illuminant discounting’.

## 1. Introduction

Color is useful for identifying objects, yet the light that reaches the eye from a surface confounds the reflectance of the surface with the spectrum of the illumination. A change of scene illumination can substantially alter the light reflected from an object, so for surface color to be useful the illuminant influence needs to be discounted. Color constancy is the visual ability to maintain a stable perception of surface color despite changes in lighting. One influential way to conceptualize a process that would achieve constancy is that the visual system estimates the (global) illumination in a scene and discounts its influence across the whole visual field, and a large body of work has sought cues in the retinal image that could support such a process (for review see [1–4]).

Two simplifications limit how far this view generalizes to natural viewing. The first concerns surface properties. Experimental stimuli have most often been flat, matte surfaces, whose diffuse reflection carries only the product of the illumination spectrum and the surface reflectance. Objects in the real world additionally elicit a percept of gloss through specular reflection, a direct mirror-like reflection of the incident light. Because specular reflection carries direct information about the illumination, the visual system might in principle use it to correct for the influence of illumination and infer surface color.

The second simplification concerns the illumination. Most studies have used a single-spectrum illuminant, ignoring the directional dependence of natural lighting: an object receives light from every direction, and the spectral content of that light varies from one incident angle to another. Outdoors, the upper part of an object may be lit directly by sunlight while light reaching its lower part is dominated by inter-reflections from the ground and nearby objects [5]. Such directional variation can even make surfaces that are indistinguishable under a single illuminant discriminable by their color [6]. Only a limited number of constancy studies have considered spatially or directionally varying illumination [7–11], even as interest in gloss, shape, material and color under natural lighting has grown [12–17]. Where color and gloss constancy have been measured under naturalistic, image-based illumination, both are found to be imperfect and to depend on the particular lighting environment [18]. We therefore still know relatively little about the extent to which color constancy holds under complex, directionally structured lighting.

Directionally dependent illumination can be simulated with a computer-graphics technique called environmental illumination [19]. A directional illumination map, defined for a particular environment, stores the light arriving from every direction toward a single point in the scene; the map allows an object to be rendered as if it were placed at that location. Typical maps contain on the order of 1024 (elevation) × 2048 (azimuth) pixels, where each pixel is a sample of light from a single direction. Rendering scenes with environmental illumination maps can capture the interaction between directionally dependent illumination and surface properties.

Given this image-formation process, one might argue that the visual system would need to deal with more than two million illuminations to achieve constancy. However, not all of these have independent effects on the light reflected from a matte object, for two reasons. First, the spatial variation of natural illumination is relatively smooth, so neighboring directions are far from independent [5,20]. Second, when light strikes a matte surface, diffuse reflection spatially averages the incident light, so by the time reflected light reaches the eye the effective angular variation of the illumination is much reduced. Even so, the influence of illumination still varies across an object’s surface, and for surfaces that additionally exhibit specular reflection the specular component carries a distorted image of the surround whose content may vary widely, with imaging geometry further modulating the relative contributions of diffuse and specular reflection.

Inverting the full illumination map would be computationally too demanding, because the proximal image confounds scene geometry (object position, illumination location, viewpoint) with the properties of surface and illumination [21]. The extent to which observers can nonetheless form stable representations of surface color across complex lighting environments may reveal the strategies that the visual system relies on. The present study therefore includes models that test how far a global, single-spectrum illuminant correction accounts for constancy under complex illumination. We stress that this modeling is not intended to rule out more sophisticated constancy mechanisms; it provides a framework within which to ask how well even a deliberately simple strategy can account for behavior, and what remains to be explained.

The choice of task also matters. Environmental illumination introduces spatial chromatic variation across an object’s surface, so classical appearance-based measures — achromatic settings [22–24] or asymmetric matching [25–29] — can be difficult when the object’s appearance itself varies spatially [30]. A measure closer to everyday behavior is selecting an object by its color, as when picking a ripe fruit from a pile. This has motivated a line of selection-based constancy experiments in which observers choose the same object across illuminations, implicitly compensating for the lighting [31–37]. These experiments, too, have used matte stimuli under single illuminants, so our understanding of constancy remains restricted to a small corner of the objects and illuminations we encounter daily. More recent studies have moved toward natural complexity, asking observers to separate illumination changes from changes in surface reflectance in hyperspectral images of natural scenes and in rendered three-dimensional scenes [38–40], probing which color shifts observers expect across illumination changes [41], and showing that natural fluctuations of daylight can reduce the number of surfaces identifiable by their color [42]. How such judgments operate when the light reaching an object varies with direction, however, remains largely unexplored.

Here we measured selection-based color constancy under real environmental illumination, and asked how far an observer that estimates one illuminant statistic and applies a diagonal correction to trichromatic sensor signals (L, M, S) can account for the behavior. On each trial four rendered objects were presented, the left and right pairs under different environments. Three objects shared a diffuse reflectance and the fourth, the target, differed; observers found the target in an “odd-one-out” task. Experiment 1 varied specularity and the presence of a surrounding context. Experiment 2 used environments with high directional anisotropy and viewing angles chosen so that the global color impression of the surround either was, or was not, representative of the light on the object.

## 2. General methods

### 2.1 Apparatus

All experiments were run in a dark room. Stimuli were shown on a calibrated CRT monitor (21 in, 1024 × 768 pixels; FP2141SB, NEC, Tokyo, Japan) driven through a visual stimulus generator (ViSaGe MkII, Cambridge Research Systems, Rochester, UK) with 14-bit intensity resolution per RGB phosphor. Gamma correction and spectral calibration were performed with a ColorCAL MKII colorimeter and a SpectroCAL MKII spectroradiometer (both Cambridge Research Systems Ltd.). Observers viewed the stimuli binocularly at a viewing distance of 92 cm, maintained with a chin rest; at this distance the test objects subtended approximately 3° of visual angle in diameter.

### 2.2 Rendering

All stimuli were rendered with computer-graphics techniques. Scene geometry — the viewpoint and the positions of objects — was defined in Blender (2.78, Blender Foundation), and images were generated by the physically based renderer Mitsuba (mitsuba-renderer.org). Mutual reflections (inter-reflections) between the objects were included in the rendering. Rendering was multispectral, from 400 to 700 nm in 10-nm steps, using the Ward reflectance model [43], in which diffuse and specular reflection sum linearly. The resulting multispectral images were converted to monitor RGB values that give the equivalent cone excitations defined by the Stockman–Sharpe cone fundamentals [44]. We used RenderToolbox4 [45] and MATLAB (2016b, MathWorks) to automate the rendering pipeline.

### 2.3 Environmental illumination

Four environmental illuminations were chosen from a publicly available database: “Distant Evening Sun (Hallstatt)” (Environment 1); “Overcast day at Techgate Donaucity” (Environment 2); “Direct Evening Sun (Schoenbrunn Vienna)” (Environment 3); and “Building Site Interior (Metro Vienna)” (Environment 4) [46]. Figure 1 shows the four maps and the chromatic distribution of all their pixels in the MacLeod–Boynton chromaticity diagram [47], computed with the Stockman–Sharpe fundamentals. For Experiment 1 we chose Environments 1 and 2, which differ in mean chromaticity and which we used previously [17]; for Experiment 2 we chose Environments 3 and 4, which have high directional anisotropy. The three-channel maps were promoted internally by Mitsuba to 31 wavelength channels [48].

**Figure 1.**
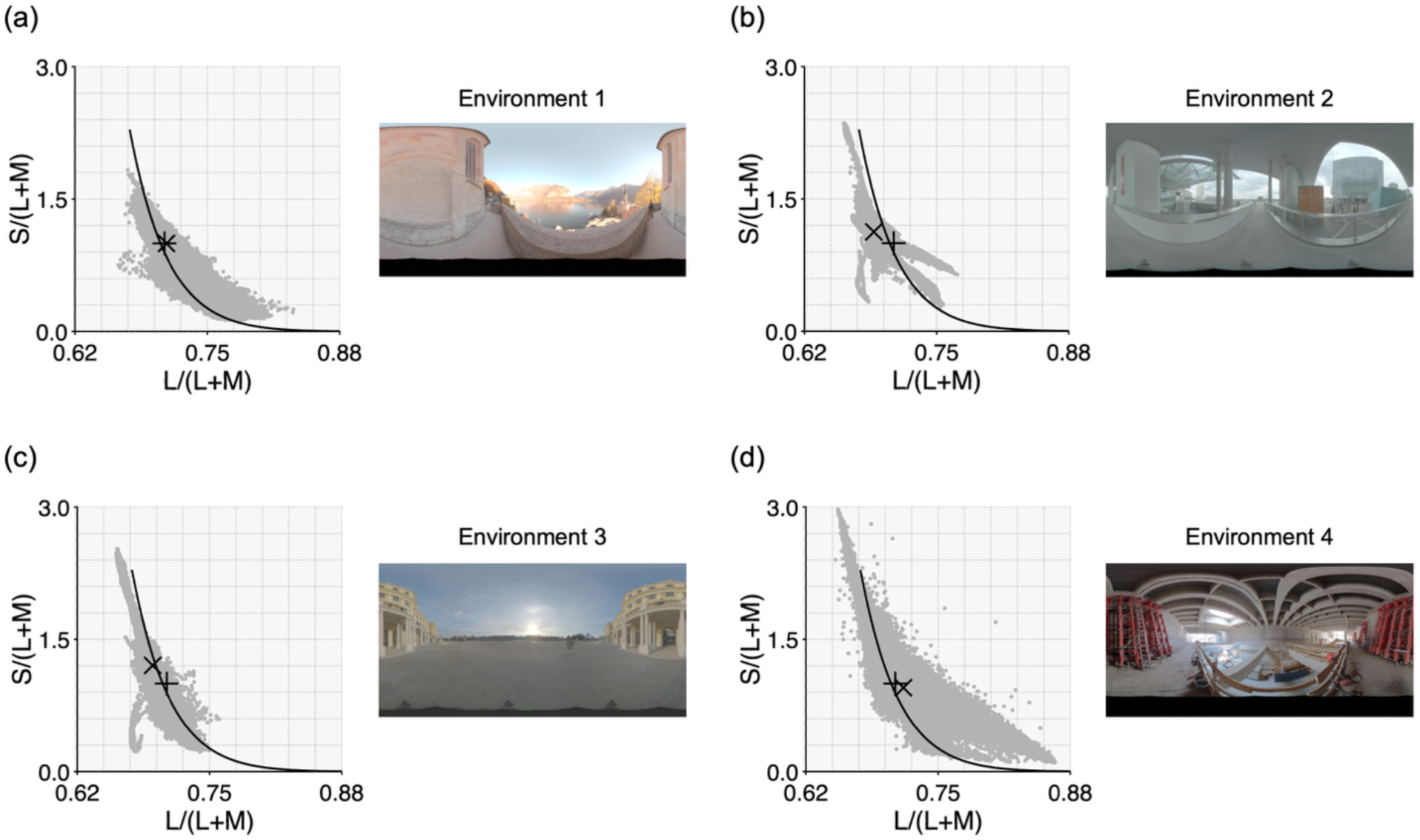
The four environmental illumination maps and the chromaticity distributions of their pixels in the MacLeod–Boynton chromaticity diagram. In each plot, the black curve indicates the black-body locus, while the plus sign and cross indicate the chromaticities of equal-energy white and the mean chromaticity across all pixels, respectively. Environments 1 and 2 (a, b) were used in Experiment 1, and Environments 3 and 4 (c, d) were used in Experiment 2.

### 2.4 Object shape and specularity

We used a bumpy sphere in Experiment 1 and a smoother, irregular object in Experiment 2. The bumpy object produces a distorted specular pattern, whereas the smoother object preserves a clearer specular image of the surround; we reasoned that the clearer image might make specular information easier to use. Two levels of specularity were used in both experiments: a fully matte surface, and a glossy surface with a specular reflectance of 0.20 at all wavelengths in the Ward model. This value was chosen so that the sum of diffuse and specular reflectance never exceeds unity at any wavelength, respecting energy conservation.

### 2.5 Surface diffuse reflectance

The choice of diffuse reflectances is critical, because it directly sets the difficulty of the task: reflectances chosen at random might be too similar for observers to discriminate, making the task impossible, and perceptual discriminability is hard to define a priori for complex stimuli. We therefore drew on our previous study [17], in which we measured discrimination thresholds for eight hue directions around equal-energy white under a single environmental illumination. Reflectances were originally selected from natural-object databases [49,50] and mixed with a flat reflectance to control the chromatic distance from equal-energy white. Discrimination was worst along the chromatic direction in which the environment holds most of its variation — near the daylight or black-body locus as shown in Figure 1. For Experiment 1 we chose, for each environment, nine reflectances whose chromaticities (when rendered under equal-energy white) lay at 2.5 times threshold from equal-energy white (Figure 2). For Experiment 2, where we had no previously measured thresholds for the new environments, we used five reflectances derived from the average of the Experiment-1 sets (flat, 0°, 90°, 180°, 270°). Figure 3 shows the rendered objects.

**Figure 2.**
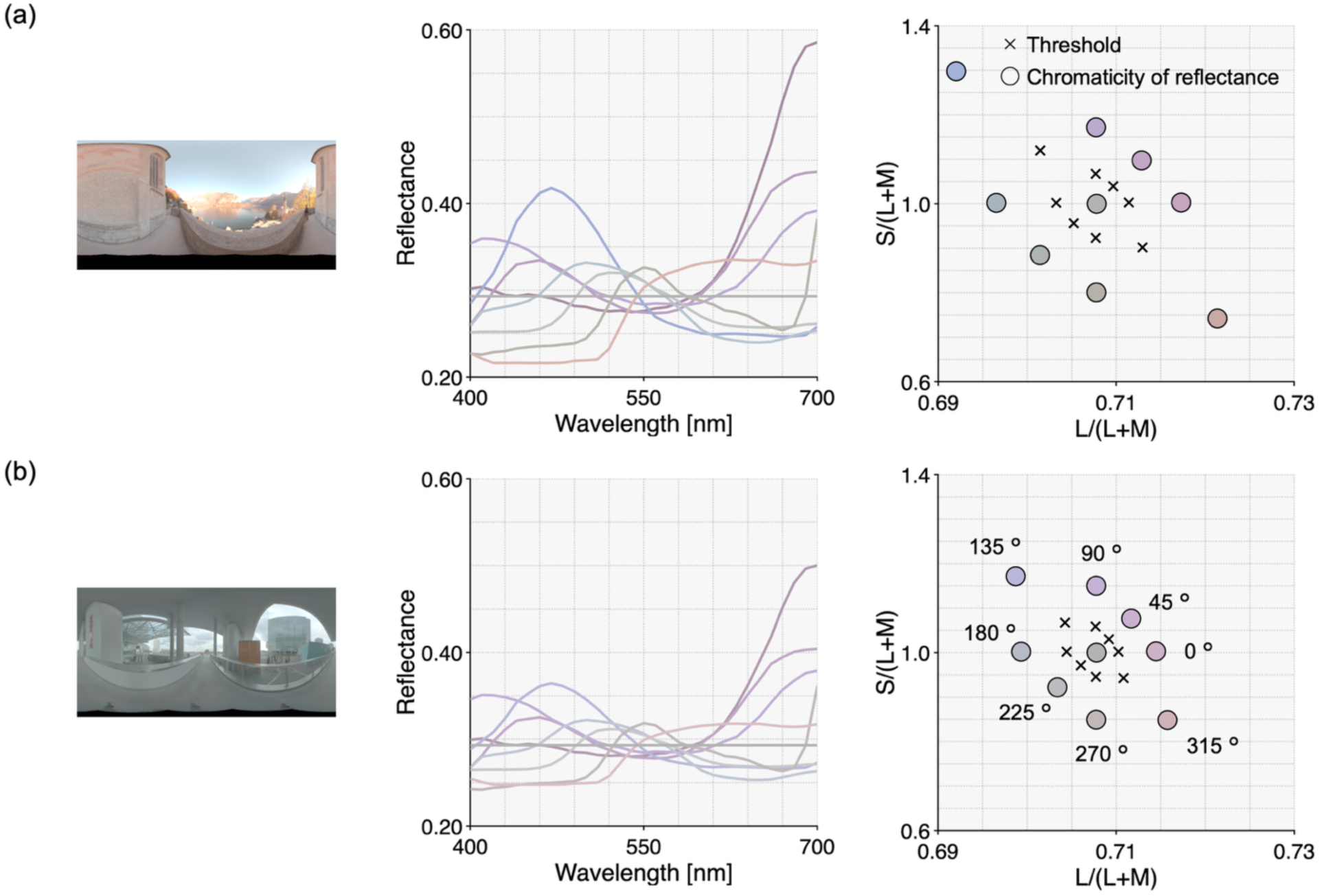
Environmental illumination (left), selected nine surface reflectances (center), and the chromatic distribution of the selected reflectances rendered under equal-energy white (right). Color-coding approximates the resultant chromaticity; black crosses show the chromatic thresholds measured previously [17]. For Environments 3 and 4, for which no thresholds were available, we used the average of the thresholds measured in Environments 1 and 2.

**Figure 3.**
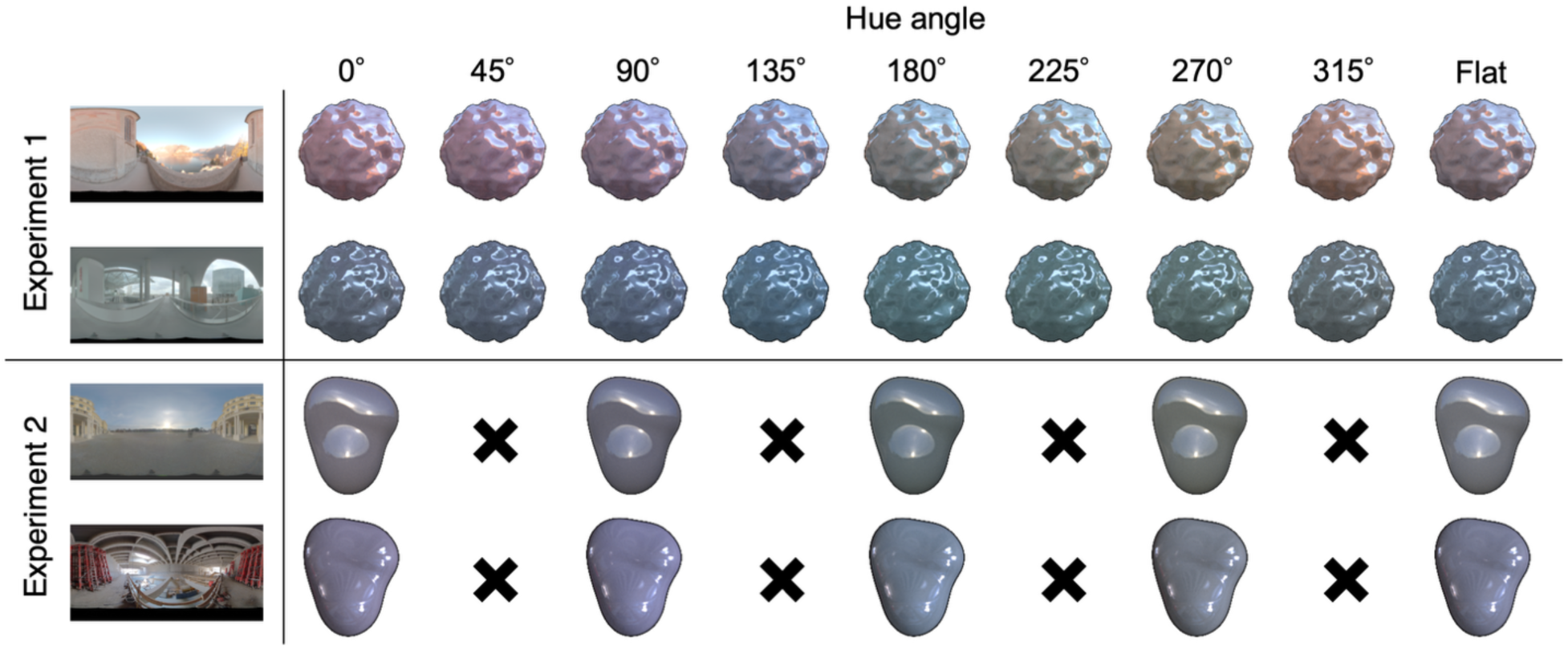
All stimuli used in the two experiments. A bumpy sphere was used for Experiment 1 and a smoother irregular object for Experiment 2. Nine reflectances were used in Experiment 1 — eight hue directions spaced 45° apart plus a neutral — and five in Experiment 2.

### 2.6 Observers and ethics

Seven observers (AKH, JH, LY, MS, SR, TD and TM; TM is an author), aged 20–32, participated in both experiments. All had corrected visual acuity and normal color vision as assessed by the HRR pseudoisochromatic plates. The design is within-observer and each observer contributed 2,304 trials in Experiment 1, so the analyses below rest on many trials per observer rather than on many observers; we return to the limits this imposes in the Discussion. The study was approved in 2016 by the Medical Sciences Inter-divisional Research Ethics Committee at the University of Oxford, in accordance with the Declaration of Helsinki.

### 2.7 Procedure

Before each block, observers adapted to 20-Hz random-dot temporal chromatic noise whose mean chromaticity and luminance equaled the mean of the nine reflectances rendered under equal-energy white [17]. On each trial, four objects were presented for 3 s (Figure 4). The left pair was rendered under one environment and the right pair under the other, separated by a vertical gray bar at the chromaticity of equal-energy white and at the mean luminance of the whole screen. Because incident light in the maps is treated as arriving from infinitely distant points, the spatial separation of the two objects within an environment has negligible effect on the light they receive. On each trial two of the reflectances were chosen; one was assigned to the target object and the other to the three distractors. Observers reported which object was the target.

**Figure 4.**
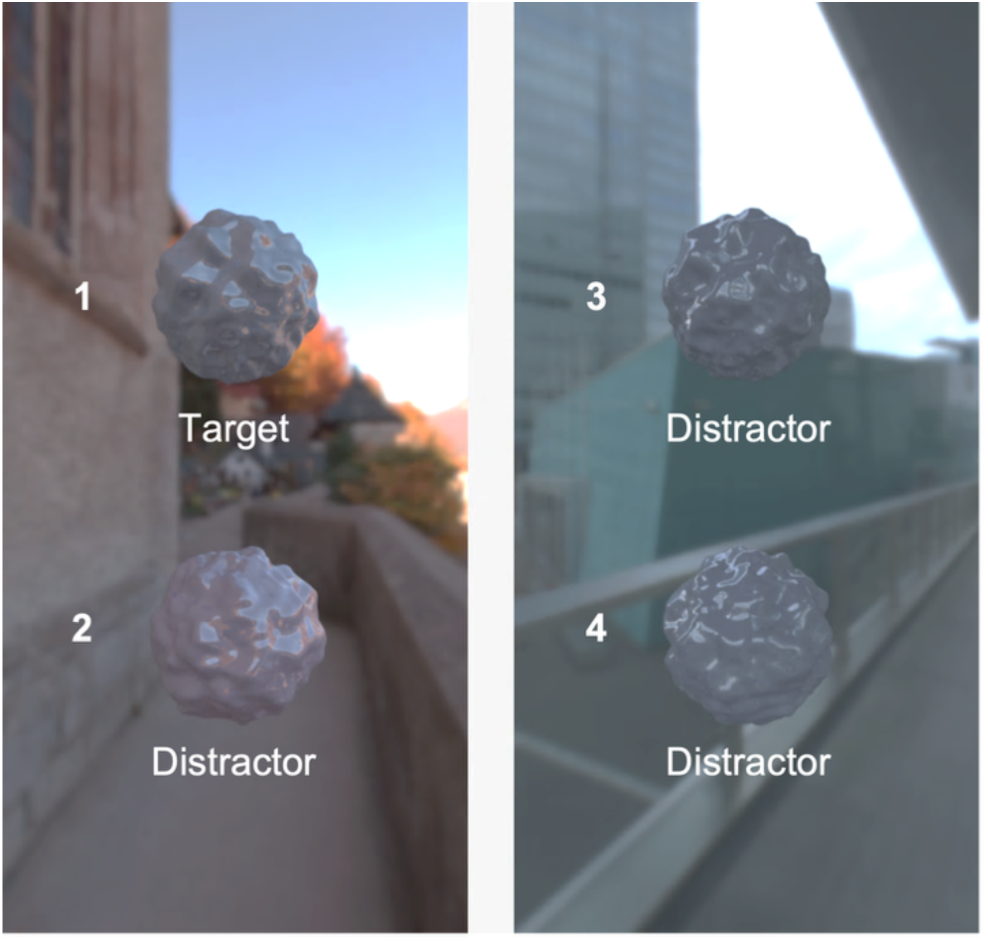
Stimulus presentation on one trial. The left and right object pairs are rendered under different environmental illuminations. Three objects share a diffuse reflectance (distractors) while the fourth (target) differs. The observer identifies the target.

The design separates two abilities. Consider a trial that has just begun. The first thing an observer notices is that the two objects on one side differ in color, so that one of them must be the target; identifying the correct side in this way depends on chromatic discrimination within a lighting environment. Choosing which of those two objects matches the pair under the other environment then requires compensating for the illumination — that is, color constancy across environments. If an observer chooses an object on the side that contains two distractors, it indicates a failure of discrimination rather than of constancy. We therefore analyzed only trials on which observers chose the correct side, so that the chance level is 50% and any shortfall reflects imperfect constancy rather than the limits of discrimination. This design is analogous to that of previous studies [35,36].

### 2.8 Computational observer models

We constructed computational observers that perform the task in the way shown in Figure 5. The models extract “uncorrected” cone signals from each object, then use an estimate of the illuminant to “correct” these signals and recover illuminant-independent surface colors. They implement a global illumination correction using parameters whose selection is informed by the long history of research with single illuminants, and so indicate what can be achieved under that simplifying assumption. The models run through every trial and choose the odd object, just as observers did. We use them to ask two questions: whether a correction of this kind can perform the task under environmental illumination at all, and whether it behaves as observers do when the cue it depends on is unreliable.

**Figure 5.**
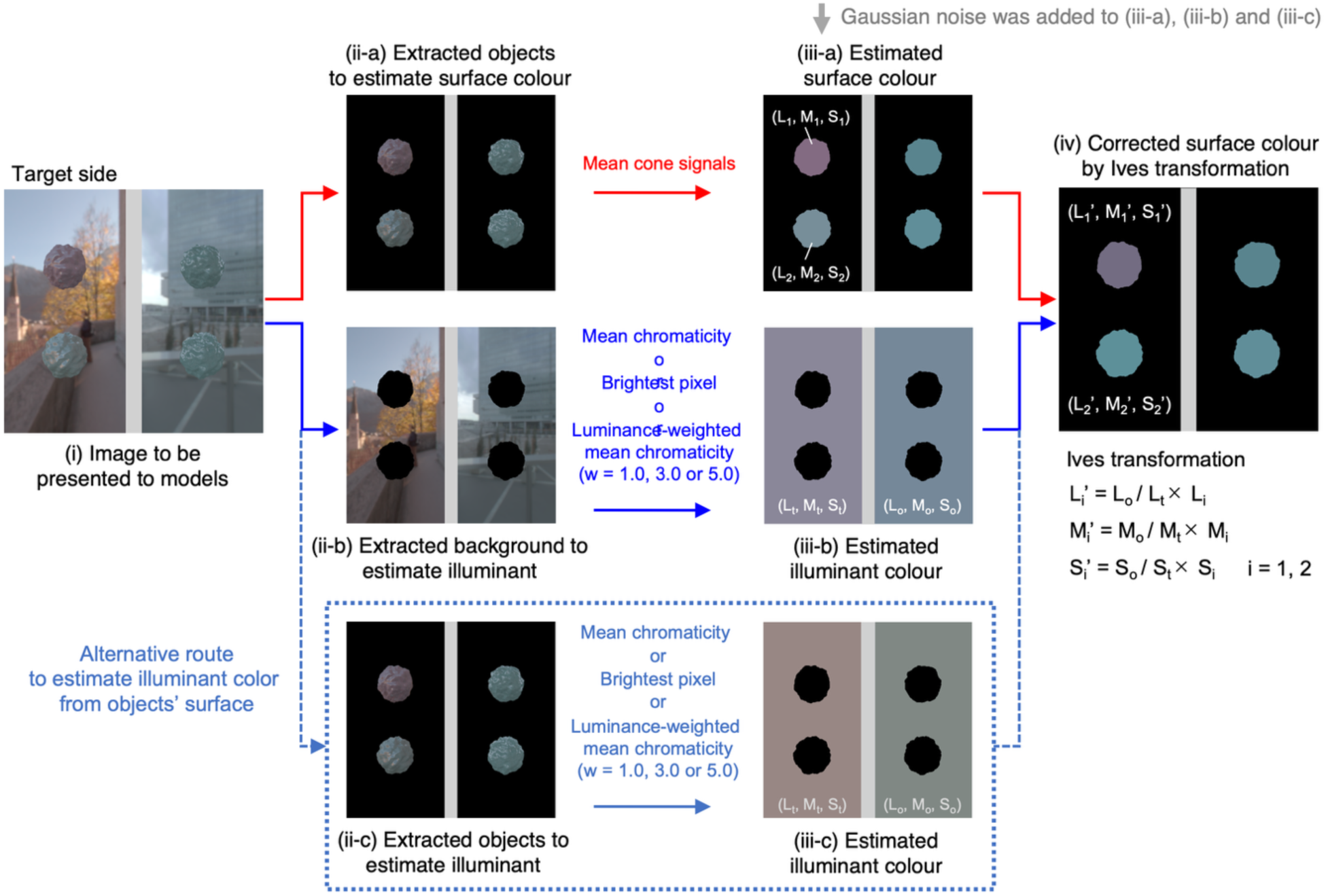
Schematic of the computational observer models. From the presented image (i) the model segments objects (ii-a) and background (ii-b); it estimates each object’s uncorrected surface color from its mean cone signals (iii-a) and the illuminant color from the background (iii-b), adding Gaussian noise to the estimates. Using the Ives transformation it corrects surface color (iv) and chooses the odd object as the chromatic outlier. An alternative route estimates the illuminant from the object regions (ii-c, iii-c).

The model first segments the image into object and surround regions, and estimates the surface color of each object separately from the illumination. Each object’s uncorrected surface color is the spatial mean of its cone signals [51]. The illuminant is estimated from the surround by one of three statistics identified in previous studies: mean chromaticity [52], the brightest pixel [53], and luminance-weighted mean chromaticity [54]. For the last, chromaticities were weighted by luminance raised to the power 1, 3 or 5; power 0 recovers mean chromaticity and power infinity the brightest pixel, so this family interpolates between the mean-chromaticity and brightest-pixel models. Because observers might, in the absence of context, estimate the illumination from an object’s own surface — particularly from specular highlights — we also built models that estimate the illuminant from the object regions. When estimating illuminant and surface colors, Gaussian noise was added independently to each estimated cone signal, with a standard deviation of 25% of that across all pixels within the environment, consistent with chromatic-discrimination data for similar stimuli [17].

The illuminant was discounted with a diagonal (Ives) transformation [55], applied to the two objects on the side containing the target so as to map the target-side illuminant onto the reference (opposite) illuminant. Writing [L, M, S] for the raw cone signals of an object and [L′, M′, S′] for the corrected signals, and with subscripts t and r for the estimated target and reference illuminants,

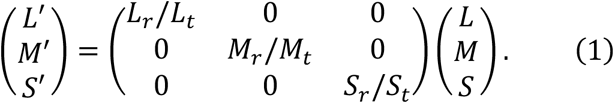

This transformation reflects the observer’s inferred judgment: having chosen the side that contains the target, the observer judges which of its two objects would match the pair under the opposite environment. No correction was applied to objects under the reference environment. The model then selected the object whose corrected chromaticity was most different from the other three in a MacLeod–Boynton diagram scaled by discrimination thresholds (0.0071 along the L/(L+M) axis and 0.1325 along the S/(L+M) axis). Selecting the object with the most different cone signals, rather than chromaticity, generally gave worse performance, presumably because of contamination by variations in intensity. A *no-correction* model performed the task without discounting the illumination, providing a baseline that quantified the magnitude of the chromatic change introduced by the illuminant difference.

## 3. Experiment 1

### 3.1 Conditions and viewing angles

We crossed two specular levels (matte, glossy) with the presence or absence of surrounding context, giving four conditions run as separate blocks. Four sessions were completed, so each observer completed 2,304 trials (144 × 4 conditions × 4 sessions).

For each of Environments 1 and 2 we selected 20 viewing angles (Figure 6). Two objects were first rendered from 36 angles (0–350° in 10° steps); depending on the angle, a different part of the environment was visible. Because environmental illumination has a high dynamic luminance range, normalizing by the single highest-luminance pixel made the images too dark, so we instead scaled each image so that 5% of pixels were saturated in at least one channel, rounding saturated values to unity. Some angles still gave dark objects, so we discarded the 16 angles with the lowest mean object luminance, leaving 20 angles per environment. Across the selected images, mean object luminance always exceeded 10 cd/m² in both matte and glossy conditions, and any saturated pixels fell in the surround rather than on the objects.

**Figure 6.**
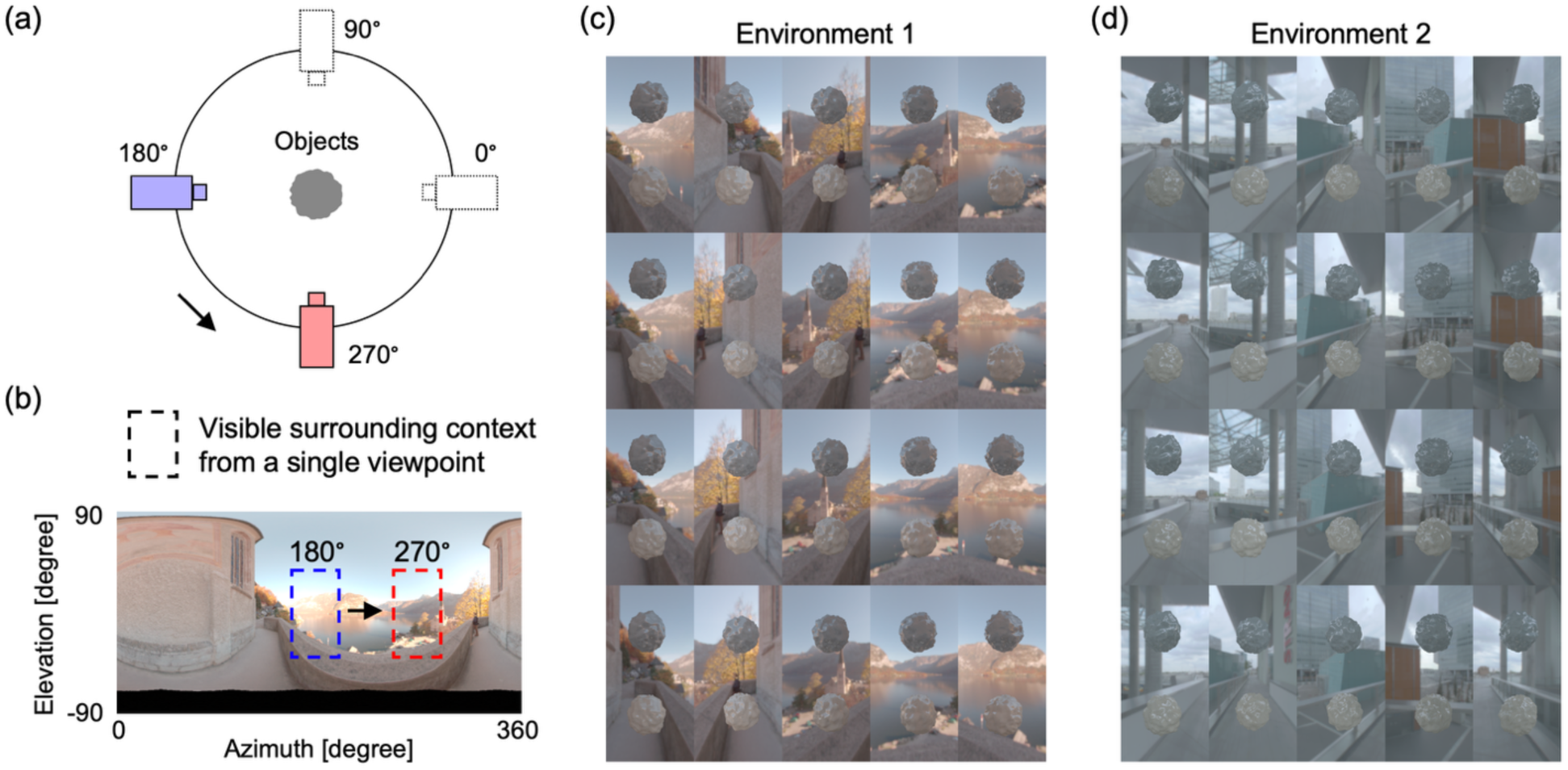
(a) The scene geometry from above. (b) The region of the environmental illumination visible from a single viewpoint. (c, d) The 20 selected viewing angles for Environments 1 and 2.

### 3.2 Results

#### 3.2.1 Human performance

Figure 7a shows percentage-correct performance across observers, with individual data shown beneath the mean. Performance was well above the 50% chance level for every observer in every condition, reaching 81.5% for matte objects with context. The no-correction model performed worse than the observers, providing a baseline measure of task difficulty arising from the magnitude of the chromatic differences introduced by the change in lighting environment.

**Figure 7.**
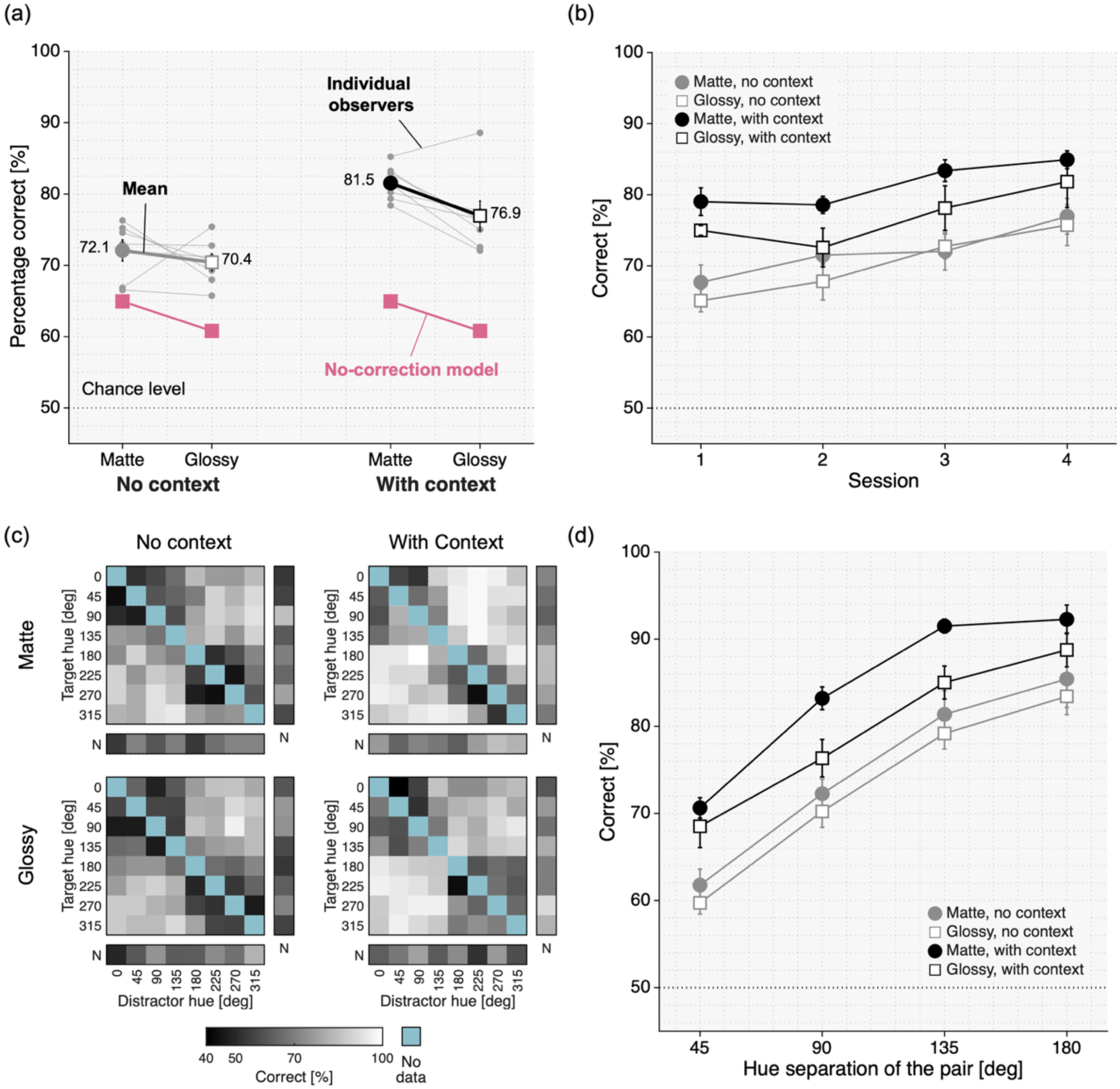
Results of Experiment 1. (a) Bold points and lines show the mean across observers; gray points and lines show individual observers. Matte and glossy conditions are grouped by context. Pink squares show the no-correction model, which performs the task without compensating for illumination; observers outperformed it in every condition. (b) Percentage correct across the four sessions for each condition (mean ± SE across observers). (c) Percentage correct for each reflectance pair, pooled across seven observers, one matrix per condition. Rows and columns are the eight hue directions, in degrees; the neutral reflectance (N) occupies its own row and column separated by a gap, because the hue circle does not continue into the neutral. Pairs near the diagonal have similar hues and were most difficult. The lower-left and upper-right triangles show trials in which the target appeared under Environments 1 and 2, respectively. Cells run from black (40%) to white (100%); no cell fell significantly below the 50% chance level after correction for multiple comparisons (one-tailed binomial test per cell, Benjamini–Hochberg FDR, *q* = 0.05). Pale blue cells contain no data. (d) Percentage correct as a function of the hue-angle separation of the pair for each condition (mean ± SE across observers). The parallel curves indicate that context improved performance equally at every separation. Horizontal dotted lines in (a), (b) and (d) mark the 50% chance level.

Observers chose the correct side on 94.9% of all trials. Out of the 576 trials per observer in each condition, the averaged side-incorrect trial number was 21.1 for the matte condition and 20.6 for the glossy condition without context, and 40.0 and 36.7 for the same two conditions with context. Side errors were about twice as frequent when context was present. Presumably, the surrounding image reduces the salience of the objects, making the within-environment discrimination harder.

A two-way repeated-measures analysis of variance (ANOVA) over the seven observers (Figure 7a) revealed a significant main effect of surrounding context (*F*(1,6) = 32.9, *p* = 0.0012) but not of specular reflection (*F*(1,6) = 3.3, *p* = 0.12); the interaction did not reach significance (*F*(1,6) = 5.3, *p* = 0.060).

Observers nonetheless performed well above chance in the no-context condition, on trials whose single image carries limited information about the illumination. We compared them with the no-correction model. Observers out-performed this model for matte objects (72.1% against 64.9%, *t*(6) = 4.81, *p* = 0.0030) and for glossy ones (70.4% against 60.8%, *t*(6) = 8.05, *p* < 0.001). Because each environment remained on the same side throughout the experiment, observers may have accumulated knowledge of the two environments across trials. The time course of performance was consistent with this possibility: across the four sessions, accuracy without context increased from 66.4% to 76.3% (Figure 7b; main effect of session, *F*(3,18) = 6.00, *p* = 0.005). Performance with context also improved, from 77.0% to 83.4% over the four sessions, and the two trends did not differ reliably (session × context interaction, *F*(3,18) = 0.90, *p* = 0.46), so this evidence alone is not conclusive: a general practice effect cannot be ruled out. Note, however, that even when context is present, a single image reveals the environment from one viewpoint only, so knowledge of the lighting environments as a whole, accumulated across trials, should raise performance in the with-context condition too. Overall, the emergence of these advantages over sessions is consistent with observers learning the two environments over time.

Figure 7c shows the performance for each reflectance-pair: one matrix per condition, in which each cell is one reflectance pair and the two triangles separate trials in which the target appeared under Environment 1 (below diagonal) from those in which it appeared under Environment 2 (above diagonal).

Two things are visible. First, difficulty is organized around the diagonal as expected — pairs of similar hue were hard, well-separated pairs easy — and second, the presence of context appears to lighten the whole matrix.

Both are tested quantitatively. Performance rose monotonically with the hue-angle separation of the pair, from 61.8% at 45° to 85.4% at 180° for matte objects without context (Figure 7d; main effect of separation *F*(3,18) = 172.6, *p* < 0.001). Context raised performance at every separation (main effect of condition *F*(3,18) = 12.0, *p* < 0.001), and it did so by the same amount at each: the four curves in Figure 7d are parallel and the interaction was flat (*F*(9,54) = 0.80, *p* = 0.61). Pairs involving the neutral reflectance were relatively difficult (65.0% and 64.9% for matte and glossy objects without context, and 73.1% and 70.3% with context).

Context also rescued pairs that were otherwise unresolvable. We tested each cell against the 50% chance level with a right-tailed one-sample *t*-test on the per-observer percentages, and corrected for the 72 comparisons within each condition with the Benjamini–Hochberg procedure at a false-discovery rate of 0.05; cells that did not pass were counted as unresolved. For matte objects, 19 of the 72 pairs were unresolved without context and 8 with context; for glossy objects, 29 without and 15 with.

The matrices in Figure 7c also show a coarser block structure: pairs whose hues both lay within 0–135° or both within 180–315° (the two diagonal quadrants of the 8 × 8 hue matrix) were more difficult than pairs that spanned the two ranges (the two off-diagonal quadrants). Pooling the trials of each quadrant type for each observer, accuracy for within-range and across-range pairs was 60.5% and 83.5% for matte and 57.5% and 82.1% for glossy objects without context, and 71.6% and 92.0% for matte and 64.9% and 88.0% for glossy objects with context. The difference was significant in every condition (paired *t*-tests across observers, *t*(6) ≥ 7.53, all *p* < 0.002 after Bonferroni correction for the four conditions) and persisted when the hue separation was matched: for pairs 45° apart, accuracy averaged over the four conditions was 58.6% within a range and 81.9% across ranges (*t*(6) = 9.91, *p* < 0.001). Accuracy therefore depended not only on the size of the hue difference but also on its direction. The 135° and 315° reflectances lie close to the daylight locus (Figure 2), so pairs within either range tend to differ along this locus, whereas pairs spanning the two ranges tend to differ orthogonally to it. Because the chromatic variation of the environments, like that of natural illumination in general, is elongated along the daylight locus (Figure 1), a surface difference in this direction is more easily confused with a change in illumination, even though all reflectances were placed at the same multiple of discrimination threshold from equal-energy white. This interpretation accords with evidence that illumination discrimination is poorer for changes along the daylight locus, particularly toward blue [4,56], and that deep neural networks trained for color constancy on naturalistic scenes are most constant for illuminations along this locus [57].

We additionally asked whether the same reflectance pair was harder when the target appeared under one environment than under the other — the comparison between the two triangles of Figure 7c. It was not: averaged over pairs, performance under the two environments differed by at most 2.8 percentage points in any condition, and none of the differences was significant (paired *t*-tests across observers, all *p* ≥ 0.075).

The next question concerned the cues and strategies observers used, which we investigated using computational models.

#### 3.2.2 Model performance

As shown in Figure 8a, with context available, the mean-chromaticity model reached 81.9% correct for matte objects and 77.5% for glossy objects, against 81.5% and 76.9% for the observers, and the luminance-weighted mean (w=1) reached 86.2% and 80.5%. Other luminance-weighted mean models (w=3 and w=5) show a similar trend. The exception was the brightest-pixel model, which fell to 52.9% (matte) and 52.1% (glossy) with context: the brightest pixel of an environmental illumination map is a small, bright, often strongly colored region such as a light source or a sky patch, and it poorly represents the light on the objects.

**Figure 8.**
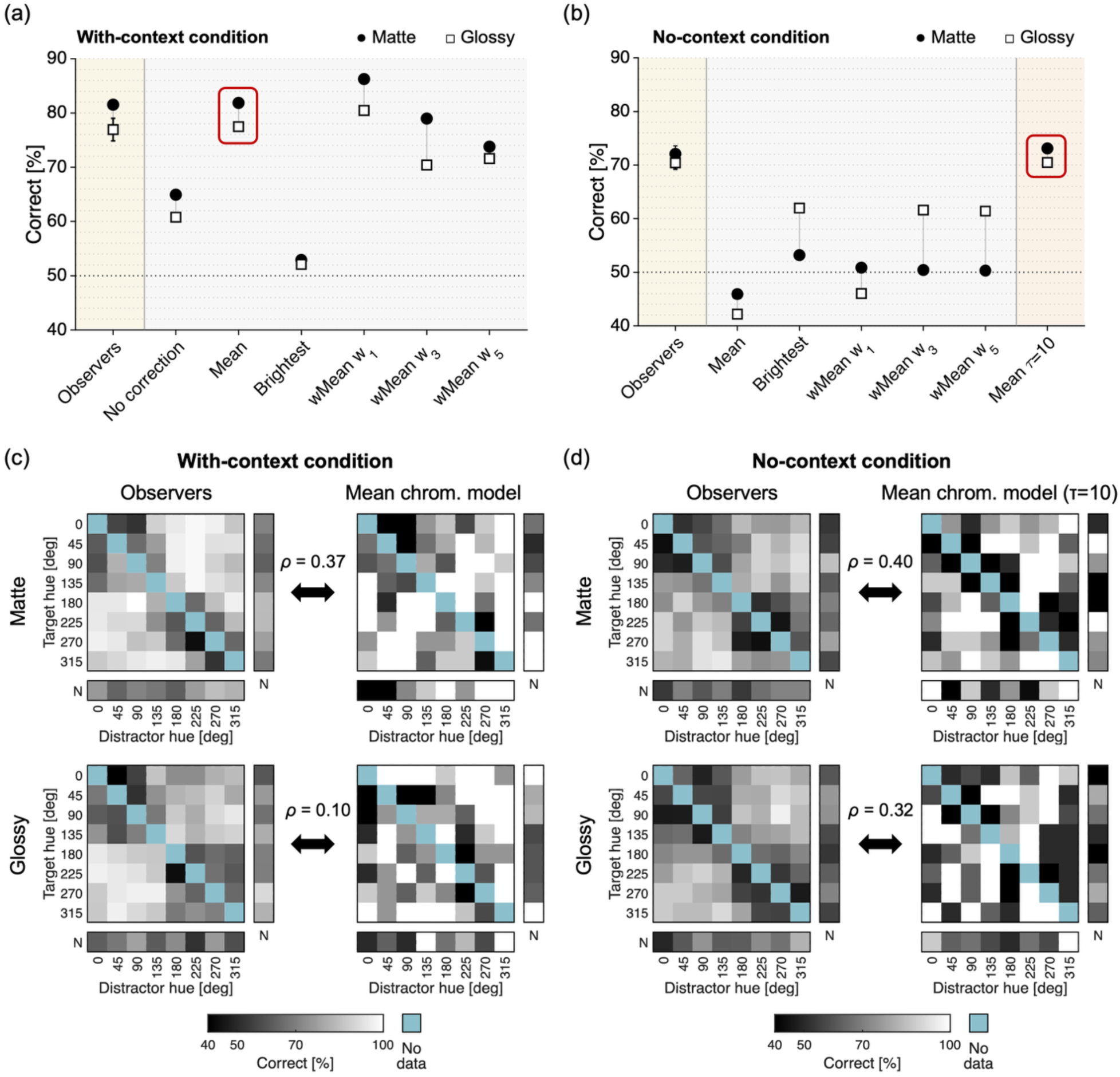
Model accuracy in Experiment 1. (a) Performance of models that estimated the illuminant from the surround, shown for matte (filled symbols) and glossy (open symbols) conditions, alongside mean observer performance at the far left. (b) The same analyses applied to the object regions, for the no-context condition. The rightmost column, separated by a vertical line, shows a different class of model: the mean-chromaticity model that integrated surround statistics from previous trials (τ = 10). Dotted horizontal lines indicate the 50% chance level. In (a) and (b), the red frame marks the model whose overall accuracy came closest to the observers’ in that condition; the matrices below are shown for these two models. (c) Reflectance-pair matrices of the observers (left) and of the mean-chromaticity model (right) in the with-context condition, for matte (top) and glossy (bottom) objects, with the Spearman rank correlation ρ between the two matrices computed over the 72 pair cells. (d) The same for the no-context condition and the temporal-integration model (Mean, τ = 10). Format and color scale as in Figure 7c.

The specular image carries usable information about the illumination. Estimating the illuminant from the object regions rather than the surround is the route available when no context is shown. As shown in Figure 8b, gloss improved the performance of statistics that focused on the brightest regions of the object. From matte to glossy objects, accuracy increased by 8.8 percentage points for the brightest-pixel model (from 53.2% to 62.0%) and by 11.2 and 11.1 points for the luminance-weighted means (w=3 and w=5), respectively. By contrast, accuracy decreased from matte to glossy objects by 3.8 points for the mean-chromaticity model and by 4.8 points for the luminance-weighted mean (w=1). Thus, gloss provided a benefit only when the weighting was sufficiently concentrated, as expected. Observers, however, showed no benefit from gloss (leftmost column). The relevant information was physically available and could be recovered by an appropriate statistic, but observers did not appear to use it.

Although some models reached human-level overall accuracy in the with-context condition, all models performed worse than observers in the no-context condition. This suggests that observers may have used additional information when no context was available. We therefore tested whether they could have accumulated information about the chromatic statistics of the two environments across trials, given that only two environments were used and each remained on the same side throughout the experiment. Models that integrated surround statistics from previous trials using an exponential weighting kernel with time constant τ became increasingly accurate in the no-context condition as τ increased: the mean-chromaticity model rose from 62.1% correct at τ = 0 [trial] to 89.5% at τ = 1000 [trial]. At τ = 10 [trial], it reached 73.1% with matte and 70.5% with glossy objects, closely matching the observers’ 72.1% and 70.4% in the same conditions (Figure 8b, rightmost column). Accumulating chromatic information across trials was therefore sufficient to support the observed level of performance and remains a plausible explanation for why observers outperformed the no-correction model when no context was present.

Finally, matching an observer’s average accuracy is not the same as behaving like that observer. To ask whether the models also succeed and fail where observers succeed and fail, we analyzed patterns of performance, for each context condition, for the model whose overall accuracy came closest to the observers’: with context, the mean-chromaticity model (red frame in Figure 8a), and without one, the temporal-integration mean-chromaticity model (τ = 10; red frame in Figure 8b). Figure 8c and d compare their reflectance-pair matrices with the observers’, through Spearman’s correlation ρ computed over the 72 pair cells. The correlation between observers sets the upper limit for these values: correlating each observer’s matrix with the pooled matrix of the remaining six gave, on average, ρ = 0.69 for matte and 0.63 for glossy objects in the with-context condition, and 0.60 and 0.58 in the no-context condition. The models fell well short of this limit: with context, the mean-chromaticity model reached ρ = 0.37 (matte) and 0.10 (glossy); without context, the τ = 10 model reached ρ = 0.40 and 0.32. Thus, even the models that best matched the observers’ overall accuracy showed less pair-by-pair agreement with observers than observers showed with one another, suggesting that the models and observers relied on different strategies. It is also worth noting that, because both observers and models solved the task well, there are too few errors to characterize their structure.

Nevertheless, Experiment 1 showed that surrounding context helped. In a Mondrian experiment a single global illuminant falls on both the test patch and its surround, so the surround is directly informative about the light on the test. Under environmental illumination, the light reflected from an object toward the eye can differ markedly from the part of the surround visible behind it. Experiment 2 exploited this dissociation by using anisotropic illuminant maps that can make the surround uninformative about the light on the object within a single trial.

## 4. Experiment 2

### 4.1 Conditions and viewing-angle selection

We crossed two specular levels (matte and glossy), the presence or absence of surrounding context, and congruency (congruent and incongruent), yielding eight conditions. Smooth objects were used so that the specular image of the environment was clearer than for the bumpy objects. Each environment was rendered from one of two viewing angles. On congruent trials, the viewing angle was selected so that the mean cone signals from the illumination on the object matched those of the surround; on incongruent trials, they did not.

Except for TM, observers were not informed of the manipulation and congruent and incongruent trials were randomly interleaved within each session. Five reflectances were used, with all pairwise combinations (_5_*C*_2_ = 10 trials) tested using the same counterbalancing as before.

Each block therefore contained 80 trials: 40 congruent and 40 incongruent. The four combinations of specular level and surrounding context were tested in separate blocks. Each observer completed four blocks for each combination, giving 1,280 trials in total, evenly divided between congruent and incongruent trials.

To select the viewing angles, we rendered a mirror sphere at the test location for every camera angle and recorded the luminance-weighted mean (w=1) of its MacLeod–Boynton chromaticities as an estimate of the light our approximately spherical objects would reflect toward the camera. We based this selection on the luminance-weighted mean because it was the most accurate of the statistics based on surrounding context in Experiment 1 (Figure 8a); the incongruent angles therefore confront the account based on surrounding-context statistics in its strongest form. We then compared this with the same quantity computed over the surround, expressed the difference in units of chromatic discrimination threshold, and chose the two angles that maximized or minimized its root-mean-square value (Figure 9). For congruent angles the object and surround means overlap in the MacLeod–Boynton diagram, whereas for incongruent angles they differ strongly. Figure 9e and 9f show example presentations: in the incongruent case the mean surround color is strongly biased — bluish where clear sky is visible, reddish where red clutter is behind the objects — and thus very different from the mean color across the objects. We asked whether observers can still perform the task when the surround’s mean cone signals are not informative about the light on the object.

**Figure 9.**
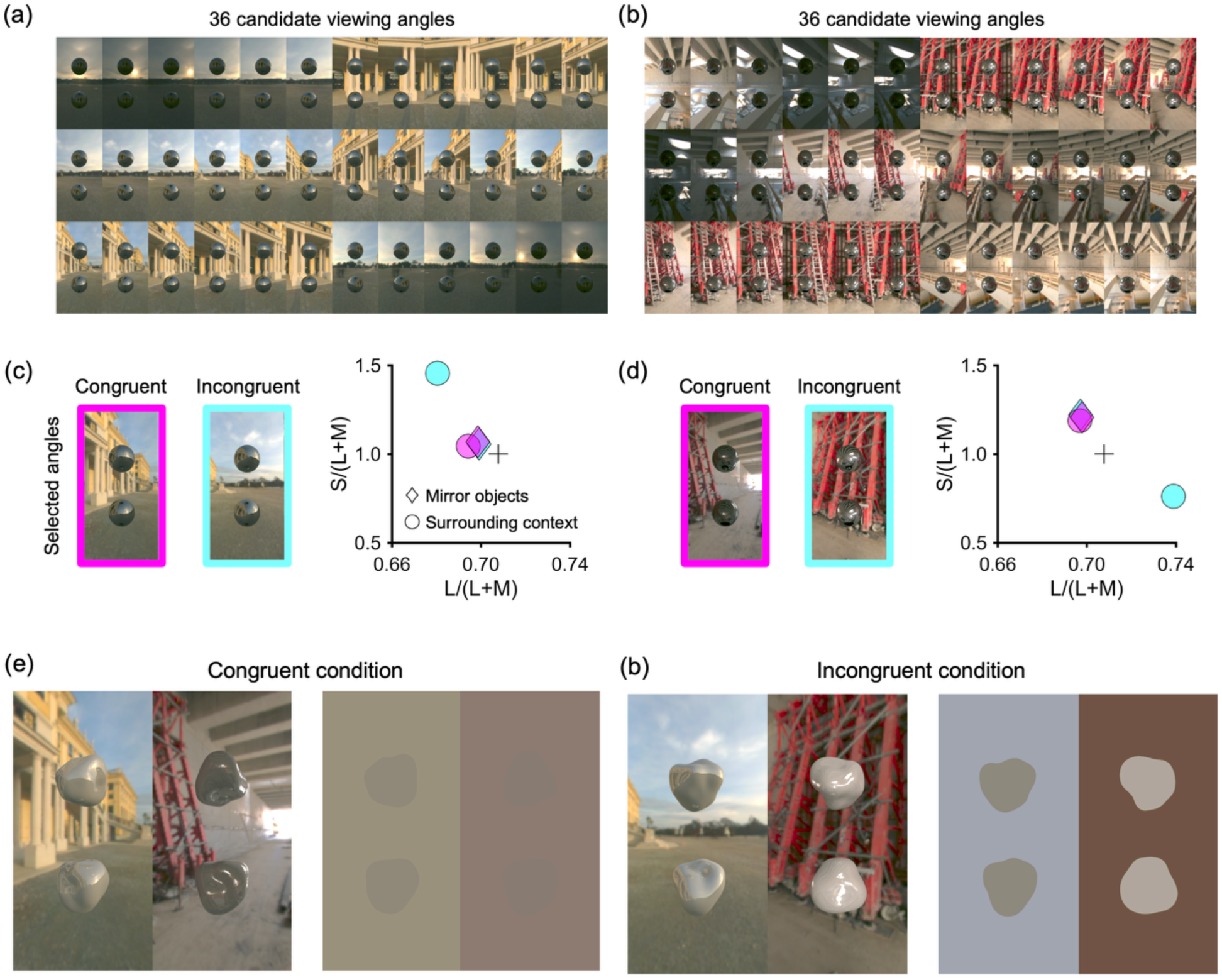
Selecting congruent and incongruent viewing angles for Experiment 2. (a, b) Environments 3 and 4 with mirror spheres rendered at all 36 candidate angles. (c, d) The two selected angles that maximize (incongruent, cyan) and minimize (congruent, magenta) the difference between the luminance-weighted mean chromaticities of the mirror objects (diamonds) and of the surround (circles), shown in the MacLeod–Boynton diagram. The difference between the diamonds across (c) and (d) is the mean illumination difference for which a constant observer should correct. Example presentations in Experiment 2 and the corresponding mean cone signals of object and surround regions for (e) a congruent condition, where the surround’s mean cone signals match the light on the objects and for (f) an incongruent condition, where they differ. In each panel the right-hand image fills the object and surround regions with the sRGB color of their mean cone signals.

### 4.2 Results

#### 4.2.1 Human performance

The central result of Experiment 2 was that the congruency manipulation imposed only a small cost on observer performance, far smaller than predicted by the no-correction model. With a surrounding context shown, making the surround uninformative about the light on the object reduced accuracy from 93.2% to 87.0% for matte objects and from 89.9% to 85.7% for glossy objects; without context, accuracy fell from 86.8% to 81.4% for matte objects and did not drop for glossy objects (82.3% to 84.6%). No condition lost more than 6.1 percentage points (Figure 10a). The no-correction model, which never consults the surrounding context and so measures what the change of viewing angles costs on its own, lost far more: 25.7 points for matte and 10.6 points for glossy objects (pink squares in Figure 10a). Whatever observers were doing, it did not depend on a single illuminant estimate read off the surround.

**Figure 10.**
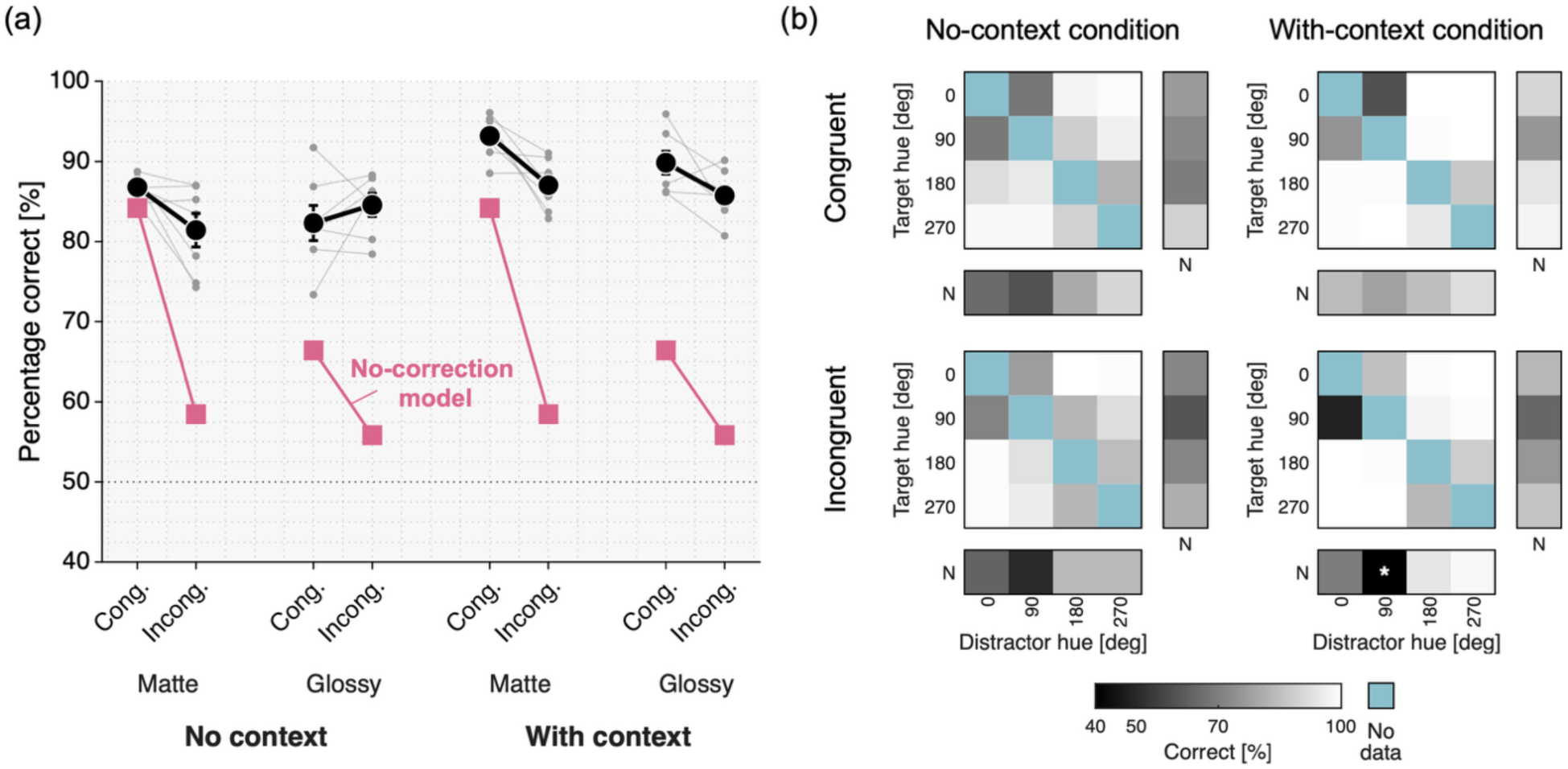
Experiment 2 human performance. (a) Percentage correct on congruent (Cong.) and incongruent (Incong.) trials in each specularity × context condition. Bold points show the mean across observers (± 1 SE); gray points and lines show individual observers. Pink squares show the no-correction model, which performs the task without compensating for illumination; because this model never consults the surround, the change from congruent to incongruent trials measures the cost of the change of viewing angles alone. The dotted line marks the 50% chance level. (b) Percentage correct for each reflectance pair, pooled over the seven observers and the two gloss levels, for the four context × congruency combinations. Format and color scale as in Figure 7c: cells run from black (40%) to white (100%), and pale blue cells contain no data. The asterisk marks the one cell significantly below the 50% chance level after correction for multiple comparisons (one-tailed binomial test per cell, Benjamini–Hochberg FDR, *q* = 0.05).

As in Experiment 1, trials on which observers chose the wrong side were excluded before scoring. Observers chose the correct side on 92.2% of trials, comparable to the 94.9% of Experiment 1, leaving 8,259 of the 8,960 trials for analysis. The slightly higher exclusion rate is consistent with the reflectances here being scaled by the average of the thresholds measured in Environments 1 and 2 [17], since no prior thresholds were available for Environments 3 and 4 (Figure 2).

A three-way repeated-measures ANOVA showed significant main effects of specularity (*F*(1,6) = 10.8, *p* = 0.017), surrounding context (*F*(1,6) = 47.1, *p* = 0.0005) and congruency (*F*(1,6) = 6.5, *p* = 0.043), and a significant specularity × congruency interaction (*F*(1,6) = 6.9, *p* = 0.039); no other interaction was significant. The simple effect of congruency was significant for matte objects (congruent > incongruent, *p* = 0.011) but not for glossy objects, and the simple effect of specularity was significant in the congruent condition (matte > glossy, *p* = 0.017) but not in the incongruent condition. Thus surrounding context helped even when it was incongruent; matte objects gave better constancy than glossy objects in the congruent condition, but this glossy disadvantage disappeared in the incongruent condition — potentially because observers exploited the specular image to infer the light on the object when the surround was misleading.

Figure 10b resolves the same performance by reflectance pair, pooled over the seven observers and the two gloss levels. The difficulty structure of Experiment 1 carried over: pairs of adjacent hues and pairs involving the neutral reflectance were hardest, and context raised performance throughout. Cells significantly below the 50% chance level were identified (one-tailed binomial test per cell on the pooled counts, Benjamini–Hochberg false-discovery rate, *q* = 0.05, within the 20 pair cells of each matrix). One cell survived the correction, showing a systematic confusion specific to the misleading surround.

#### 4.2.2 Model performance

We applied the same models to Experiment 2 (Figure 11a). On congruent trials the models did well, and the best of them came close to the observers: the luminance-weighted mean (w=1) reached 92.8% against 91.5% for observers, with the other statistics between 53.6% and 69.1%. For incongruent trials, the models that estimate an illuminant from the surround performed far worse. The luminance-weighted mean (w=1) fell by 44.2 points, from 92.8% to 48.6%, below the 50% chance level; the luminance-weighted mean (w=3) fell 22.3 points to 46.8%, and the mean-chromaticity model fell 15.9 points to 47.5%. The brightest-pixel model is the one surround-based model that lost nothing (53.6% to 53.9%), but it did not solve the task well in the first place.

**Figure 11.**
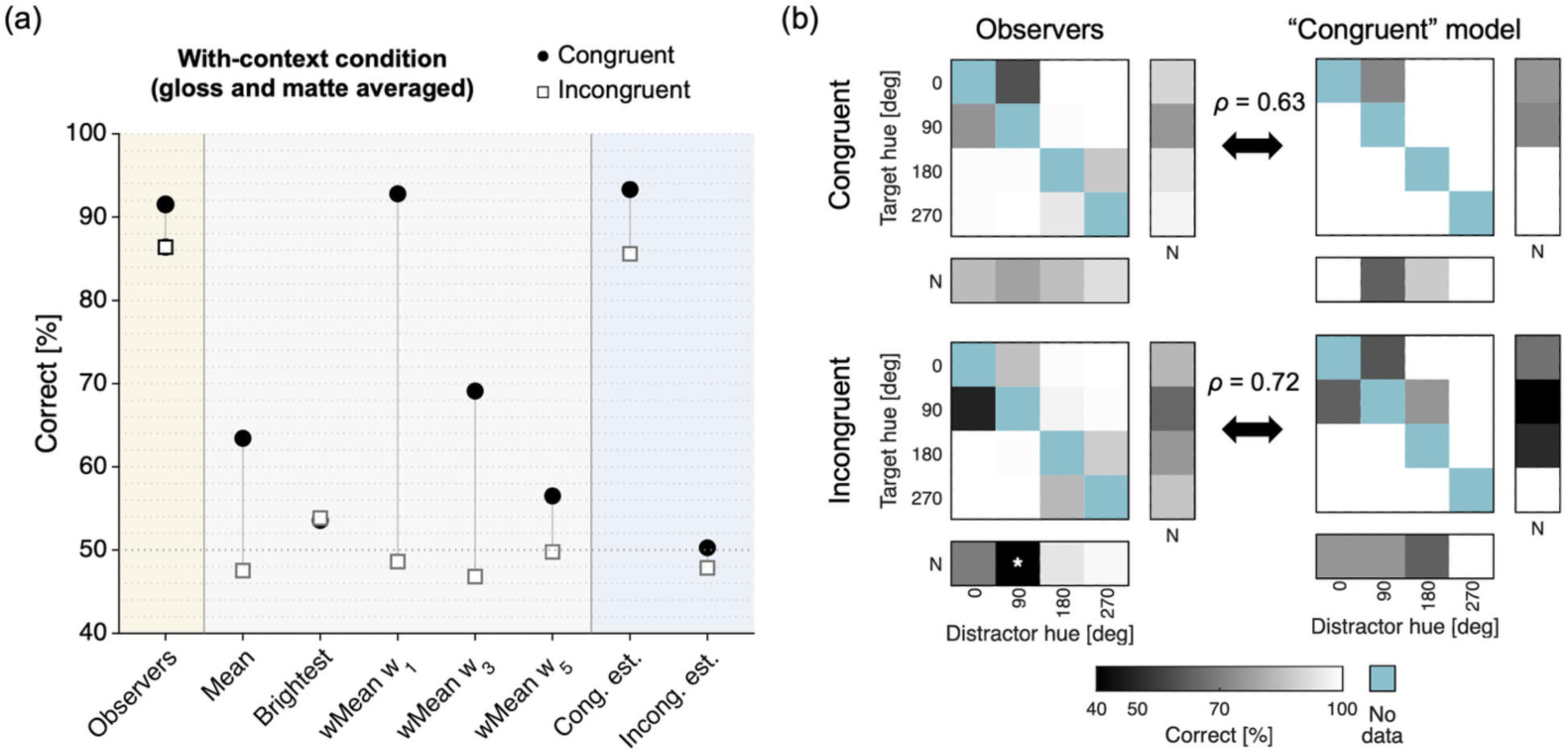
Experiment 2 model accuracy, restricted to the two conditions with context and averaged over matte and glossy. (a) Percentage correct on congruent (filled circles) and incongruent (open squares) trials; background tints separate the observers (mean ± 1 SE across the seven observers; yellow) from the surround-based models, and those in turn from the two models that hold a fixed estimate (blue). Mean, Brightest and wMean w1–w5 estimate the illuminant from the surround of the current trial and apply a diagonal correction; the model that best matched the observers on congruent trials, the luminance-weighted mean (w=1), fell furthest on incongruent trials, below the 50% chance level (dotted line). Cong. est. and Incong. est. instead hold a fixed estimate — the luminance-weighted mean (w=1) of the congruent or of the incongruent surround, respectively — and divide it out on every trial. The no-correction baseline is shown in Figure 10a. (b) Reflectance-pair performance matrices of the observers (left; the with-context matrices of Figure 10b) and of the congruent-estimate model (right), on congruent (top) and incongruent (bottom) trials, with the Spearman rank correlation between the two matrices computed over the 20 reflectance-pair cells. Format and color scale as in Figure 7c.

One alternative is that observers learned each environment’s illuminant across the session and corrected each image using this remembered estimate rather than the current surround. We tested this account by applying the temporal-integration models to Experiment 2 (not shown in Figure 11a). Indeed, their performance changed little between congruent and incongruent trials, dropping by only 2.7 percentage points for the mean-chromaticity model and 3.1 points for the luminance-weighted mean (w=1). However, performance was near chance even on congruent trials (52.3% and 52.7%, respectively), far below the observers’ 91.5%. Thus, although accumulating surround statistics was sufficient in Experiment 1, it cannot explain performance in Experiment 2, where the estimate averages across congruent and incongruent presentations of the same environment.

The two rightmost models in Figure 11a sharpen this: the failure of the temporal-integration models lies in what they average, not in relying on a stored estimate. Each of the two models holds a fixed surround estimate instead of reading the surround of the current trial — the luminance-weighted mean (w=1) of the congruent surround for the congruent-estimate model, and of the incongruent surround for the incongruent-estimate model. The congruent-estimate model behaved like the observers — 93.3% correct on congruent trials and 85.6% on incongruent ones, a loss of 7.7 points against the observers’ 5.1 — while the incongruent-estimate model remained near chance (50.3% and 47.9%). The resemblance extends to the level of reflectance pairs (Figure 11b): the congruent-estimate model’s matrix ranks the pairs much as the observers’ does, with Spearman correlations over the 20 pair cells of *ρ* = 0.63 on congruent trials (*p* = 0.003) and *ρ* = 0.72 on incongruent trials (*p* < 0.001). The correlation between observers again sets the upper limit for these values: correlating each observer’s matrix with the pooled matrix of the remaining six gave, on average, *ρ* = 0.75 on congruent and 0.86 on incongruent trials. A global correction anchored to the light actually falling on the objects reproduces the observers’ behavioral patterns, on congruent and incongruent trials alike.

## 5. Discussion

We conducted two psychophysical experiments to measure how well observers identify surface colors across environmental illuminations. Constancy was generally good and orderly. Difficulty was governed, as expected, by the chromatic separation of the two reflectances. A surrounding context raised performance by the same amount at every chromatic separation, whereas gloss brought observers no benefit.

The statistics we tested — mean chromaticity, the brightest pixel, the luminance-weighted mean — were all devised for flat Mondrian displays under a single light, and it was not obvious that they would transfer to illumination whose spectrum depends on direction. In Experiment 1, to some extent, they did: when an informative surround was available, several models approached or even exceeded observers’ overall accuracy. However, their agreement with observers was limited when responses were compared trial by trial.

Even in Experiment 2, when the surround was made uninformative about the light on the objects, observers appeared to build up knowledge of each lighting environment and to use it selectively, perhaps by treating objects and background as distinct sources of evidence rather than pooling all into one illuminant estimate — much as color judgments in immersive virtual environments depend on how the local surround and the wider illumination are combined [58,59]. For instance, an observer who has learned that the reddish regions of Environment 4 are local objects — the red clutter of the building site — rather than a property of the illumination can exclude them from the estimate of the global illuminant. Observers may also draw on expectations about how surface colors typically shift across illumination changes [41]. These results suggest a dynamic learning in real-world color constancy that has received little attention in previous research.

One consistent result is that specular highlights did not help. In principle they could: a highlight is a partial image of the light falling on the object, even though it is contaminated by the diffuse reflection, and algorithms have long exploited this to recover illuminant chromaticity [60]. The psychophysical evidence is mixed. Highlights and other specular cues have been reported to improve constancy when the scene is otherwise impoverished [61], when surface and illumination geometry can be inferred [62], and alongside other three-dimensional cues such as binocular disparity [63]; but observers often fail to exploit them when they are available [64], gain nothing from the motion of a glossy object [65], and show color and lightness judgments that are largely unaffected by gloss [14,18,30]. Our results fall on the negative side, and they add a physical control: the information was demonstrably present, since object-based statistics that concentrate on the brightest pixels gained 9 to 14 percentage points when the objects were glossy (Figure 8b). Observers, given the same images, gained little and in some conditions lost a little. Three factors may reconcile this. Our scenes were cue-rich, and highlights appear to matter chiefly when other cues are weak [61]. Under environmental illumination a highlight is not a compact bright spot but a spatially complex pattern whose chromaticity varies from point to point, so the mapping from highlight to illuminant is far less direct than under a point source [66]. We suspected that the bumpy objects of Experiment 1 blurred this pattern, but the smoother objects of Experiment 2 gave no specular benefit either. Finally, the specular component contaminates the estimate of the diffuse surface color [30], so gloss can cost something in a surface-color task even while carrying illuminant information. Information that is present in an image, information the visual system extracts, and information it uses in a given judgment are separable.

We note that what makes a model human-like is that it succeeds and fails on the same trials as observers. Comparing error patterns rather than accuracies is therefore a critical perspective [67,68]. In our case, several models matched the observers’ overall accuracy well, but their agreement with observers across individual reflectance pairs was much lower than the agreement among observers. The comparison, however, is limited by the fact that observers and models both solved the task well, so errors were few and unevenly spread over the pairs, and the error structure is correspondingly poorly resolved. Distinguishing models on this basis requires conditions in which observers make many, systematic errors. Sampling a wider range of object shapes, reflectances and lighting environments — rather than repeating a small set — would lower performance into the range where errors become diagnostic [18], and would at the same time test whether the conclusions here generalize beyond the stimulus conditions we used.

Our models test statistics we could name in advance, and if observers use image measurements we did not anticipate, no amount of fitting will reveal them. An alternative is to collect human judgments on a large and diverse stimulus set and to train image-computable models to predict those judgments, then ask what the trained model measures [69–72]. We recently took this approach to gloss perception and found that networks small enough to be interpreted — in some cases a single convolutional kernel — reproduce human judgments across many shapes and lighting environments, and the kernels reveal which image structures observers rely on [73]. The same recipe suits the present problem, which is harder than gloss in that the relevant information is chromatic and distributed across object and surround. It would demand human data on a scale well beyond seven observers, but that is now within reach, and it offers a way to identify the computations observers actually perform rather than the ones we can articulate.

Several limitations are noted. Each environment was presented on the same side throughout, so the information observers used when no context was available may have been learned environment identity rather than anything estimated from the current image; our temporal-integration models show that accumulating chromatic statistics across trials is sufficient to solve Experiment 1, but an experiment that varied the side assignment and increased the number of environments would be needed to establish that observers did so. Constancy was also good but not complete — no observer exceeded 90% correct in Experiment 1 — and one likely reason is that observers viewed the scenes on a display rather than from within them, and so were never adapted to either environment as an observer standing in it would be. In addition, because the stimuli were presented on a flat display, the specular reflections carried no binocular disparity, a three-dimensional cue that has been shown to improve color constancy [63]; its absence may be another reason why gloss was not as helpful as it might have been.

This study joins a growing effort to measure color constancy under illumination with real-world complexity: immersive virtual-reality scenes have shown that constancy generalizes to full-field, spectrally complex environments and have begun to dissect the mechanisms that support it [58,59] and how it depends on the field of view [74]; measurements in everyday settings report near-complete constancy [75]; an equivalent-noise analysis of lightness constancy in naturalistic rendered scenes has quantified how much of observers’ performance is limited by scene-to-scene variation rather than by their own sensory noise [76]; and image-computable models are increasingly evaluated against human judgments [77,78]. Also in the present study, encouragingly, human color constancy appears more robust than any single conventional mechanism can explain, pointing instead to the accumulation and selective use of information about the lighting environment.

## Funding

T.M. was supported by a Sir Henry Wellcome Postdoctoral Fellowship from the Wellcome Trust (218657/Z/19/Z), Aso Scholarship, JASSO Oversea Scholarship, Kikawada Foundation Scholarship, and the Sasakawa Fund.

## Acknowledgments

For the purpose of open access, the author has applied a CC BY public copyright license to any Author Accepted Manuscript version arising from this submission.

## Disclosures

The authors declare no conflicts of interest.

## Data availability

The individual observer responses, along with the analysis and figure-generation code supporting the findings of this study, are openly available at https://github.com/takuma929/reflectance_identification.

## Notes

### Competing Interest Statement

The authors have declared no competing interest.

## References

1. H. E. Smithson, “Sensory, computational and cognitive components of human colour constancy,” Philosophical Transactions of the Royal Society B: Biological Sciences 360, 1329–1346 (2005).

2. A. Hurlbert, “Colour constancy,” Current Biology 17, R906–R907 (2007).

3. D. H. Foster, “Color constancy,” Vision Research 51, 674–700 (2011).

4. A. Hurlbert and C. Yu, “Seeing the light: perception and discrimination of illumination color,” Annual Review of Vision Science 11, 267–301 (2025).

5. T. Morimoto, S. Kishigami, J. M. M. Linhares, S. M. C. Nascimento, and H. E. Smithson, “Hyperspectral environmental illumination maps: characterizing directional spectral variation in natural environments,” Optics Express 27, 32277–32293 (2019).

6. T. Morimoto, J. M. M. Linhares, S. M. C. Nascimento, and H. E. Smithson, “How many surfaces can you distinguish by color? Real environmental lighting increases discriminability of surface colors,” Optics Express 32, 34246–34253 (2024).

7. J. N. Yang and S. K. Shevell, “Surface color perception under two illuminants: The second illuminant reduces color constancy,” Journal of Vision 3, 369–379 (2003).

8. H. Boyaci, K. Doerschner, and L. T. Maloney, “Perceived surface color in binocularly viewed scenes with two light sources differing in chromaticity,” Journal of Vision 4, 664–679 (2004).

9. K. Doerschner, H. Boyaci, and L. T. Maloney, “Human observers compensate for secondary illumination originating in nearby chromatic surfaces,” Journal of Vision 4, 92–105 (2004).

10. R. J. Lee and H. E. Smithson, “Context-dependent judgments of color that might allow color constancy in scenes with multiple regions of illumination,” Journal of the Optical Society of America A 29, A247–A257 (2012).

11. H. Boyaci, K. Doerschner, J. L. Snyder, and L. T. Maloney, “Surface color perception in three-dimensional scenes,” Visual Neuroscience 23, 311–321 (2006).

12. R. W. Fleming, R. O. Dror, and E. H. Adelson, “Real-world illumination and the perception of surface reflectance properties,” Journal of Vision 3, 347–368 (2003).

13. K. Doerschner, H. Boyaci, and L. T. Maloney, “Testing limits on matte surface color perception in three-dimensional scenes with complex light fields,” Vision Research 47, 3409–3423 (2007).

14. M. Olkkonen and D. H. Brainard, “Perceived glossiness and lightness under real-world illumination,” Journal of Vision 10(9), 5 (2010).

15. I. Motoyoshi and H. Matoba, “Variability in constancy of the perceived surface reflectance across different illumination statistics,” Vision Research 53, 30–39 (2012).

16. Y. Morgenstern, W. S. Geisler, and R. F. Murray, “Human vision is attuned to the diffuseness of natural light,” Journal of Vision 14(9), 15 (2014).

17. T. Morimoto and H. E. Smithson, “Discrimination of spectral reflectance under environmental illumination,” Journal of the Optical Society of America A 35, B244–B255 (2018).

18. T. Morimoto, A. Akbarinia, K. R. Storrs, J. R. Cheeseman, H. E. Smithson, K. R. Gegenfurtner, and R. W. Fleming, “Color and gloss constancy under diverse lighting environments,” Journal of Vision 23(7), 8 (2023).

19. P. Debevec, “Rendering synthetic objects into real scenes: bridging traditional and image-based graphics with global illumination and high dynamic range photography,” in Proceedings of the 25th Annual Conference on Computer Graphics and Interactive Techniques (SIGGRAPH ’98) (ACM, 1998), pp. 189–198.

20. R. O. Dror, A. S. Willsky, and E. H. Adelson, “Statistical characterization of real-world illumination,” Journal of Vision 4, 821–837 (2004).

21. R. W. Fleming, “Visual perception of materials and their properties,” Vision Research 94, 62–75 (2014).

22. D. H. Brainard, “Color constancy in the nearly natural image. 2. Achromatic loci,” Journal of the Optical Society of America A 15, 307–325 (1998).

23. M. D. Fairchild and P. Lennie, “Chromatic adaptation to natural and incandescent illuminants,” Vision Research 32, 2077–2085 (1992).

24. J. S. Werner and J. Walraven, “Effect of chromatic adaptation on the achromatic locus: The role of contrast, luminance and background color,” Vision Research 22, 929–943 (1982).

25. L. E. Arend, A. Reeves, J. Schirillo, and R. Goldstein, “Simultaneous color constancy: papers with diverse Munsell values,” Journal of the Optical Society of America A 8, 661–672 (1991).

26. K.-H. Bäuml, “Simultaneous color constancy: How surface color perception varies with the illuminant,” Vision Research 39, 1531–1550 (1999).

27. D. H. Brainard and B. A. Wandell, “Asymmetric color matching: how color appearance depends on the illuminant,” Journal of the Optical Society of America A 9, 1433–1448 (1992).

28. F. W. Cornelissen and E. Brenner, “Simultaneous colour constancy revisited: an analysis of viewing strategies,” Vision Research 35, 2431–2448 (1995).

29. L. Arend and A. Reeves, “Simultaneous color constancy,” Journal of the Optical Society of America A 3, 1743–1751 (1986).

30. B. Xiao and D. H. Brainard, “Surface gloss and color perception of 3D objects,” Visual Neuroscience 25, 371–385 (2008).

31. D. H. Brainard, N. P. Cottaris, and A. Radonjic, “The perception of colour and material in naturalistic tasks,” Interface Focus 8, 20180012 (2018).

32. Y. Ling and A. Hurlbert, “Color and size interactions in a real 3D object similarity task,” Journal of Vision 4, 721–734 (2004).

33. A. Radonjić, N. P. Cottaris, and D. H. Brainard, “Color constancy supports cross-illumination color selection,” Journal of Vision 15(6), 13 (2015).

34. A. Radonjić, N. P. Cottaris, and D. H. Brainard, “The relative contribution of color and material in object selection,” PLoS Computational Biology 15, e1006950 (2019).

35. R. Robilotto and Q. Zaidi, “Limits of lightness identification for real objects under natural viewing conditions,” Journal of Vision 4, 779–797 (2004).

36. R. Robilotto and Q. Zaidi, “Lightness identification of patterned three-dimensional, real objects,” Journal of Vision 6, 18–36 (2006).

37. Q. Zaidi and M. Bostic, “Color strategies for object identification,” Vision Research 48, 2673–2681 (2008).

38. R. Ennis and K. Doerschner, “Disentangling simultaneous changes of surface and illumination,” Vision Research 158, 173–188 (2019).

39. S. M. C. Nascimento and D. H. Foster, “Misidentifying illuminant changes in natural scenes due to failures in relational colour constancy,” Proceedings of the Royal Society B: Biological Sciences 290, 20231676 (2023).

40. L. S. Virtanen, M. Olkkonen, and T. P. Saarela, “The effect of illumination cues on color constancy in simultaneous identification of illumination and reflectance changes,” Journal of Vision 25(6), 4 (2025).

41. H. Karimipour and C. Witzel, “Colour expectations across illumination changes,” Vision Research 222, 108451 (2024).

42. D. H. Foster, “Fluctuating environmental light limits number of surfaces visually recognizable by colour,” Scientific Reports 11, 2102 (2021).

43. G. J. Ward, “Measuring and modeling anisotropic reflection,” ACM SIGGRAPH Computer Graphics 26, 265–272 (1992).

44. A. Stockman and L. T. Sharpe, “The spectral sensitivities of the middle- and long-wavelength-sensitive cones derived from measurements in observers of known genotype,” Vision Research 40, 1711–1737 (2000).

45. B. S. Heasly, N. P. Cottaris, D. P. Lichtman, B. Xiao, and D. H. Brainard, “RenderToolbox3: MATLAB tools that facilitate physically based stimulus rendering for vision research,” Journal of Vision 14(2), 6 (2014).

46. B. Vogl, “Light probes,” http://dativ.at/lightprobes/ (accessed 25 July 2026).

47. D. I. A. MacLeod and R. M. Boynton, “Chromaticity diagram showing cone excitation by stimuli of equal luminance,” Journal of the Optical Society of America 69, 1183–1186 (1979).

48. B. Smits, “An RGB-to-spectrum conversion for reflectances,” Journal of Graphics Tools 4, 11–22 (1999).

49. M. J. Vrhel, R. Gershon, and L. S. Iwan, “Measurement and Analysis of Object Reflectance Spectra,” Color Research & Application 19, 4–9 (1994).

50. L. Chittka, A. Shmida, N. Troje, and R. Menzel, “Ultraviolet as a component of flower reflections, and the colour perception of hymenoptera,” Vision Research 34, 1489–1508 (1994).

51. L. S. Virtanen, M. Olkkonen, and T. P. Saarela, “Color ensembles: Sampling and averaging spatial hue distributions,” Journal of Vision 20(5), 1 (2020).

52. G. Buchsbaum, “A spatial processor model for object colour perception,” Journal of the Franklin Institute 310, 1–26 (1980).

53. E. H. Land, “The retinex theory of color vision,” Sci. Am. 237, 108–128 (1977).

54. B.-G. Khang and Q. Zaidi, “Illuminant color perception of spectrally filtered spotlights,” Journal of Vision 4, 680–692 (2004).

55. H. E. Ives, “The Relation between the Color of the Illuminant and the Color of the illuminated Object,” Transactions of the Illuminating Engineering Society 7, 62–72 (1912).

56. S. Aston, A. Radonjić, D. H. Brainard, and A. C. Hurlbert, “Illumination discrimination for chromatically biased illuminations: implications for color constancy,” Journal of Vision 19(3), 15 (2019).

57. A. Flachot, A. Akbarinia, H. H. Schütt, R. W. Fleming, F. A. Wichmann, and K. R. Gegenfurtner, “Deep neural models for color classification and color constancy,” Journal of Vision 22(4), 17 (2022).

58. R. Gil Rodríguez, L. Hedjar, M. Toscani, D. Guarnera, G. C. Guarnera, and K. R. Gegenfurtner, “Color constancy mechanisms in virtual reality environments,” Journal of Vision 24(5), 6 (2024).

59. R. Gil Rodríguez, L. Hedjar, B. Kilic, and K. R. Gegenfurtner, “Colour constancy: colour interaction between local surround and illumination in virtual reality scenarios,” bioRxiv 2026.02.12.705534 (2026).

60. H.-C. Lee, “Method for computing the scene-illuminant chromaticity from specular highlights,” Journal of the Optical Society of America A 3, 1694–1699 (1986).

61. R. Wedge-Roberts, S. Aston, U. Beierholm, R. Kentridge, A. Hurlbert, M. Nardini, and M. Olkkonen, “Specular highlights improve color constancy when other cues are weakened,” Journal of Vision 20(12), 4 (2020).

62. R. J. Lee and H. E. Smithson, “Low levels of specularity support operational color constancy, particularly when surface and illumination geometry can be inferred,” Journal of the Optical Society of America A 33, A306 (2016).

63. J. N. Yang and S. K. Shevell, “Stereo disparity improves color constancy,” Vision Research 42, 1979–1989 (2002).

64. J. N. Yang and L. T. Maloney, “Illuminant cues in surface color perception: tests of three candidate cues,” Vision Research 41, 2581–2600 (2001).

65. R. J. Lee and H. E. Smithson, “Motion of glossy objects does not promote separation of lighting and surface colour,” Royal Society Open Science 4, 171290 (2017).

66. T. Morimoto, R. J. Lee, and H. E. Smithson, “When the brightest is not the best: illuminant estimation from the geometry of specular highlights,” Proceedings of the Royal Society B: Biological Sciences 293, 20260317 (2026).

67. F. A. Wichmann, D. H. J. Janssen, R. Geirhos, G. Aguilar, H. H. Schütt, M. Maertens, and M. Bethge, “Methods and measurements to compare men against machines,” Electronic Imaging 29(14), 36–45 (2017).

68. R. Geirhos, K. Meding, and F. A. Wichmann, “Beyond accuracy: quantifying trial-by-trial behaviour of CNNs and humans by measuring error consistency,” Advances in Neural Information Processing Systems 33, 13890–13902 (2020).

69. N. Kriegeskorte, “Deep Neural Networks: A New Framework for Modeling Biological Vision and Brain Information Processing,” Annual Review of Vision Science 1, 417–446 (2015).

70. R. M. Cichy, A. Khosla, D. Pantazis, A. Torralba, and A. Oliva, “Comparison of deep neural networks to spatio-temporal cortical dynamics of human visual object recognition reveals hierarchical correspondence,” Scientific Reports 6, 27755 (2016).

71. N. J. Majaj and D. G. Pelli, “Deep learning—Using machine learning to study biological vision,” Journal of Vision 18(13), 2 (2018).

72. S. Ponting, T. Morimoto, and H. E. Smithson, “Modeling surface color discrimination under different lighting environments using image chromatic statistics and convolutional neural networks,” Journal of the Optical Society of America A 40, A149–A159 (2023).

73. T. Morimoto, A. Akbarinia, K. R. Storrs, J. R. Cheeseman, H. E. Smithson, K. R. Gegenfurtner, and R. W. Fleming, “Human gloss perception reproduced by tiny neural networks,” Nature Human Behaviour 10, 1340–1355 (2026).

74. K. Duay and T. Nagai, “Impact of field of view on color constancy in virtual reality,” Journal of the Optical Society of America A 42, B199–B213 (2025).

75. K. R. Gegenfurtner, D. Weiss, and M. Bloj, “Color constancy in real-world settings,” Journal of Vision 24(2), 12 (2024).

76. V. Singh, J. Burge, and D. H. Brainard, “Equivalent noise characterization of human lightness constancy,” Journal of Vision 22(5), 2 (2022).

77. H. Heidari-Gorji and K. R. Gegenfurtner, “Object-based color constancy in a deep neural network,” Journal of the Optical Society of America A 40, A48–A56 (2023).

78. H. Heidari-Gorji, R. Gil Rodríguez, and K. R. Gegenfurtner, “Human-aligned evaluation of a pixel-wise DNN color constancy model,” Frontiers in Human Neuroscience 20, 1809976 (2026).

